# Hunting *Drosophila* viruses from wild populations: a novel isolation approach and characterization of viruses

**DOI:** 10.1101/2022.09.16.508214

**Authors:** Gaspar Bruner-Montero, Carlos Luque, Shuai Dominique Ding, Jonathan P. Day, Francis M. Jiggins

**Affiliations:** Department of Genetics, University of Cambridge, Cambridge CB2 3EH, UK

## Abstract

Metagenomic studies have demonstrated that viruses are extremely diverse and abundant in insects, but the difficulty of isolating them means little is known about the biology of these newly discovered viruses. To overcome this challenge in *Drosophila*, we created a cell line with increased susceptibility to infection and detected novel viruses by the presence of double-stranded RNA. We demonstrate the utility of these tools by isolating La Jolla virus (LJV) and Newfield virus (NFV) from several wild *Drosophila* populations. These viruses have different potential host ranges, with distinct abilities to replicate in five *Drosophila* species. Similarly, in some species they cause high mortality and in others they are comparatively benign. In three species, NFV but not LJV caused large declines in female fecundity. This sterilization effect was associated with differences in tissue tropism, as NFV but not LJV was able to infect *Drosophila melanogaster* ovaries. We saw a similar effect in the invasive pest of fruit crops *Drosophila suzukii*, where oral infection with NFV caused reductions in the fecundity, suggesting it has potential as a biocontrol agent. In conclusion, a simple protocol allowed us to isolate new viruses and demonstrate that viruses identified by metagenomics have a large effect on the fitness of the model organism *D. melanogaster* and related species.

## Introduction

Over recent years metagenomics has revolutionized the discovery of new viruses, revealing extraordinary viral diversity and advancing our understanding of virus evolution [1]. However, the isolation of new viruses has not kept pace with this endeavor, thanks to a lack of appropriate culture techniques, frequently resulting from the strict requirement of a living host cell for replication and the difficulty of detecting new viruses. This means that the impact of these viruses on the health of their hosts is unknown and their biology largely unstudied. This pattern is exemplified by research on *Drosophilidae*, where metagenomic approaches have allowed the discovery of over 100 viruses, including many that infect *D. melanogaster* [2–5]. However, only a handful of these viruses have subsequently been isolated and studied in the laboratory [6–8]. Most virus research in *Drosophila* relies on viruses isolated decades ago [9], sometimes from other species of insect.

A striking result from metagenomic and PCR surveys has been that the prevalence of virus infection is extremely high in many natural populations of *Drosophila* [5, 10]—in *D. melanogaster* most individuals in the wild are thought to be infected with one or more virus [5, 10, 11]. The effect of these viruses on flies is mostly unknown, but where they have been isolated these viruses commonly reduce both the survival and reproductive success of flies, suggesting they are important determinants of the fitness. Drosophila C virus (DCV), which is widely studied but rare in nature, is virulent, with infections in laboratory stocks causing the death of larvae and pupae, and flies injected with the virus dying within days [12]. Other viruses like Drosophila A virus (DAV) and the DNA viruses Drosophila innubila Nudivirus (DiNV) and Kallithea virus (KV) reduce host fecundity and kill their hosts, albeit at a slower rate than typically reported for DCV [8, 13, 14]. Some viruses, such as Galbut virus and sigma virus, have less obvious symptoms [12]. However, indirect estimates have suggested that even the sigma virus causes an ~20% reduction in the fitness of infected flies in the wild [15]. The impact of viruses on host fitness is corroborated by the observation that genes involved in defending flies against viruses commonly evolve rapidly [16], and natural selection has favored genetic variants that increase virus resistance [17, 18]. Together, these results demonstrate that viruses are common and sometimes virulent pathogens of *D. melanogaster*.

Insect pathogenic viruses are promising biological control agents. Baculoviruses, which are large double-stranded DNA viruses, are the only group that have been widely deployed commercially, mainly for the control of lepidopteran pests [19]. However, field trials have shown that entomopathogenic RNA viruses also have promise in biocontrol [20]. The large number of new viruses being discovered through metagenomics suggests that there may be many biocontrol agents awaiting discovery, with the first step being to isolate and characterize these viruses.

In this study we have developed an accessible protocol to detect and isolate a wide range of viruses of wild *Drosophila* populations. This is based on the development of a cell line with increased susceptibility to viruses. We then follow O’Brian and colleagues in using the presence of double stranded RNA (dsRNA), which is produced by many viruses during their replication, as a tool to detect novel viruses in these cells [21]. We then demonstrate the utility of this approach by isolating two *Drosophila* viruses, the iflavirus La Jolla virus (LJV) and the permutotetravirus Newfield virus (NFV), which we show have distinct impacts on the fitness of several *Drosophila* species.

## Results

### A cell culture system to isolate novel viruses

To facilitate the isolation of viruses, we created a *Drosophila* cell line where RNAi was suppressed, increasing its susceptibility to infection. To achieve this, we stably transfected DL2 cells with a plasmid expressing V5-tagged B2, a double-stranded RNA (dsRNA) binding protein produced by Flock House virus (FHV) [22], creating a cell line which we named DL2-B2. Using immunofluorescent microscopy with anti-V5 antibodies, we confirmed that these cells expressed the B2 protein in the cytoplasm after inducing expression with CuSO_4_ for 24 h.

Double-stranded RNA is produced by positive-strand RNA viruses, double-strand RNA viruses and DNA viruses [23]. We therefore used a dsRNA enzyme-linked immunosorbent assay (ELISA) based on the J2 monoclonal antibody as a tool to detect novel viruses (see also [21]). To validate this approach and confirm that the DL2-B2 cells had increased susceptibility to viruses, cells were infected with the dicistrovirus DCV at different concentrations. Consistently more dsRNA was detected in the infected cells at 48 hours post infection (hpi) than in the uninfected cells, confirming that this virus produces dsRNA (Figure 1A). Furthermore, there was more dsRNA in the cells expressing the B2 protein, indicating that this succeeded in making the cells more susceptible to viral infection. This effect was significant at all but the highest and lowest concentrations of the virus (Figure 1A; Tukey’s HSD Test comparing cell types, *p*<0.01 at all viral doses)—the decline in dsRNA levels at the highest viral concentrations in DL2-B2 cells is likely caused by the virus causing extensive cell death. In our controls, there was no significant difference in dsRNA concentration between uninfected DL2-B2 and DL2 cells (Figure 1A; Tukey’s HSD Test: *t*= 0.31, d.f. = 79, p = 0.76).

**Figure 1.**
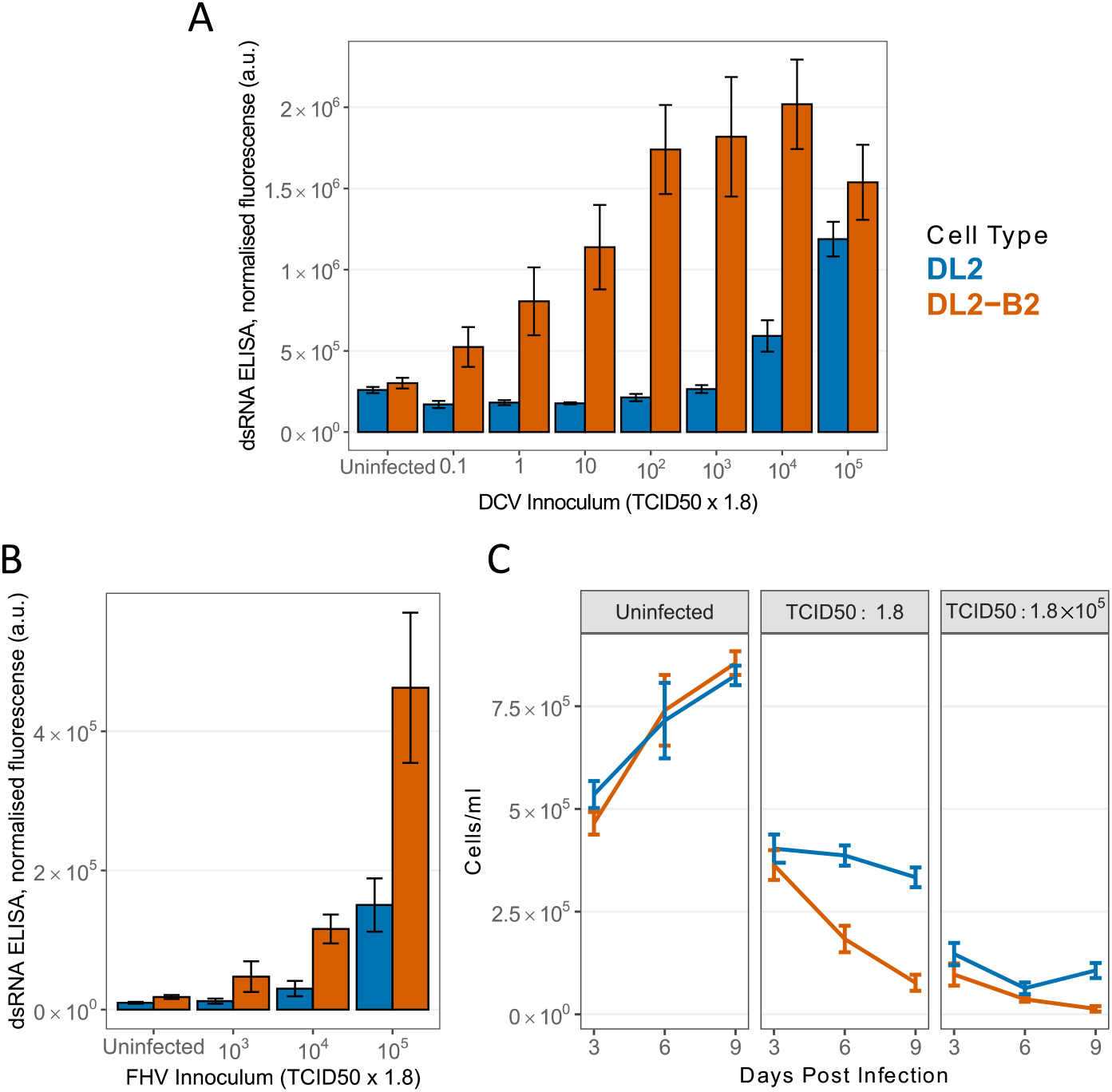
A cell culture system for isolating viruses. (A-B) DL2 cells and DL2 cells expressing the B2 suppressor of RNAi (DL2-B2) were inoculated with a range of doses of DCV and FHV, and dsRNA was detected using ELISA based on J2 monoclonal antibodies 48 hpi. The bar indicates mean fluorescence intensity (arbitrary units, a.u.) normalized to cell seeding density. (C) Cell density 3, 6 and 9 days post infection (dpi) with DCV in DL2 and DL2-B2 cells. Bars represent standard errors. TCID50 was estimated using DL2 cells.

To confirm these results with a different virus, we repeated the experiment with FHV, an alphanodavirus. Again, expressing the B2 protein in cells led to higher dsRNA levels (Figure 1B), and this effect was significant across the different inoculum concentrations (ANOVA on infected cells only, main effect cell line: *F*=11.9, d.f.=1,24, *P*=0.002). Again, there was no significant difference in dsRNA concentration between uninfected DL2-B2 and DL2 cells (Tukey’s HSD Test: *t*= 1.69, d.f. = 24, *p* = 0.11).

Many viruses also cause morphological changes to cells (cytopathic effect) and cell death, providing a second way to detect viruses. We therefore repeated the infections with DCV and counted the number of cells over a time-course. When uninfected, the density of cells increased over time, and expressing the B2 protein had no significant effect on cell growth (Figure 1C; ANOVA, main effect cell type: *F*=0.01, d.f.=1,20, *p*=0.38). When infected with a low concentration of virus, the cell density declined at a considerably faster rate in cells expressing B2 (Figure 1C; ANOVA, time × cell type: *F*=14.1, d.f.=1,32, *p*=0.0007). When infected with a high concentration of virus, cell numbers were strongly reduced in both cell lines, but again this effect was strongest when expressing B2 (Figure 1C; ANOVA, main effect cell type: *F*=11.6, d.f.=1,32, *p*=0.002). Therefore, cell death provides another indicator of viral infection, and this effect is enhanced by expressing B2.

### Isolation of novel viruses

Several species of wild-caught *Drosophila* were collected from populations across Europe and pooled in groups of 5–10 of the same sex and species. These were homogenized, filtered with a 0.22µm PVDF filter to remove microbial contamination, and then added to DL2-B2 cells. After 8–10 days, we inspected these samples under a microscope and used ELISA to test for the presence of dsRNA. We selected 19 samples that both exhibited clear cytopathic effect and had high dsRNA concentrations, and tested them for common *Drosophila* viruses using qPCR (Table S1). Nine samples were infected with La Jolla virus (LJV), two were coinfected with the DNA virus Kallithea and Newfield virus (NFV), one was coinfected with NFV and LJV, and two with DCV. As DCV is widely used in our laboratory and is uncommon in the wild [5], these two samples were discarded. We did not detect known viruses in the remaining five samples.

As these isolates sometimes consisted of a mixture of viruses, we attempted to clone single viruses using three rounds of end-point-dilution culture. In every round we performed eight replicates of 12 serial ten-fold dilutions, and selected virus from the most dilute sample that resulted in infection. At the end of this we only detected the presence of NFV in the samples that were previously coinfected with Kallithea virus or LJV.

We selected eleven samples that showed consistent CPE and used RNA sequencing (RNA-seq) to examine whether samples were infected by previously unknown viruses. To check for viral contamination, we also sequenced the DL2-B2 cell line. After assembling RNA-seq reads that did not map to the *D. melanogaster* genome or known *Drosophila* viruses, we used BLAST (Basic Local Alignment Search Tool) to identify possible viruses. Among the viral contigs with an open reading frame encoding at least 40 amino acids, the top BLAST hit was always La Jolla virus, Newfield virus, or Hubei permutotetra-like virus 3. As Hubei permutotetra-like virus 3 is a close relative of Newfield virus [24], we concluded there were no previously unknown viruses in our samples nor contaminating viruses in the DL2-B2 cell line. When mapping to known *Drosophila* viruses, four samples were infected with LJV and three with NFV. We noted a low level of LJV reads in the NFV samples, but as LJV was not detected in these samples in subsequent experiments using quantitative PCR (qPCR) we attributed this to contamination during the preparation of the libraries.

To obtain genome sequences of the newly isolated viruses, we used our sequence reads to modify a reference genome sequence of these viruses, and these were aligned with published sequences from these viruses to reconstruct their phylogenetic relationships. NFV is a positive-sense single-stranded RNA virus with a genome size of ~4.7 kb belonging to an unclassified genus within the *Permutotetraviridae* family [5]. Our three isolates of NFV were all from samples of *D. melanogaster* collected in Cambridge, UK, and they formed a clade that was related to two isolates from *D. melanogaster* from Australia (Figure 2A). LJV is an abundant virus in several species of *Drosophila* [4, 5]. It has a positive-sense single-stranded RNA genome of ~9.7 kb and belongs to an unclassified genus within the *Iflaviridae* family [5]. Our four isolates came from two pools of *D. simulans* from Cyprus, a pool of *D. melanogaster* from Cyprus, and a pool of *D. repleta* from Spain. They fell into two clades on the tree, and are closely related to sequences that were mostly obtained from *D. melanogaster* and *D. simulans* collected in Europe and America (Figure 2B).

**Figure 2.**
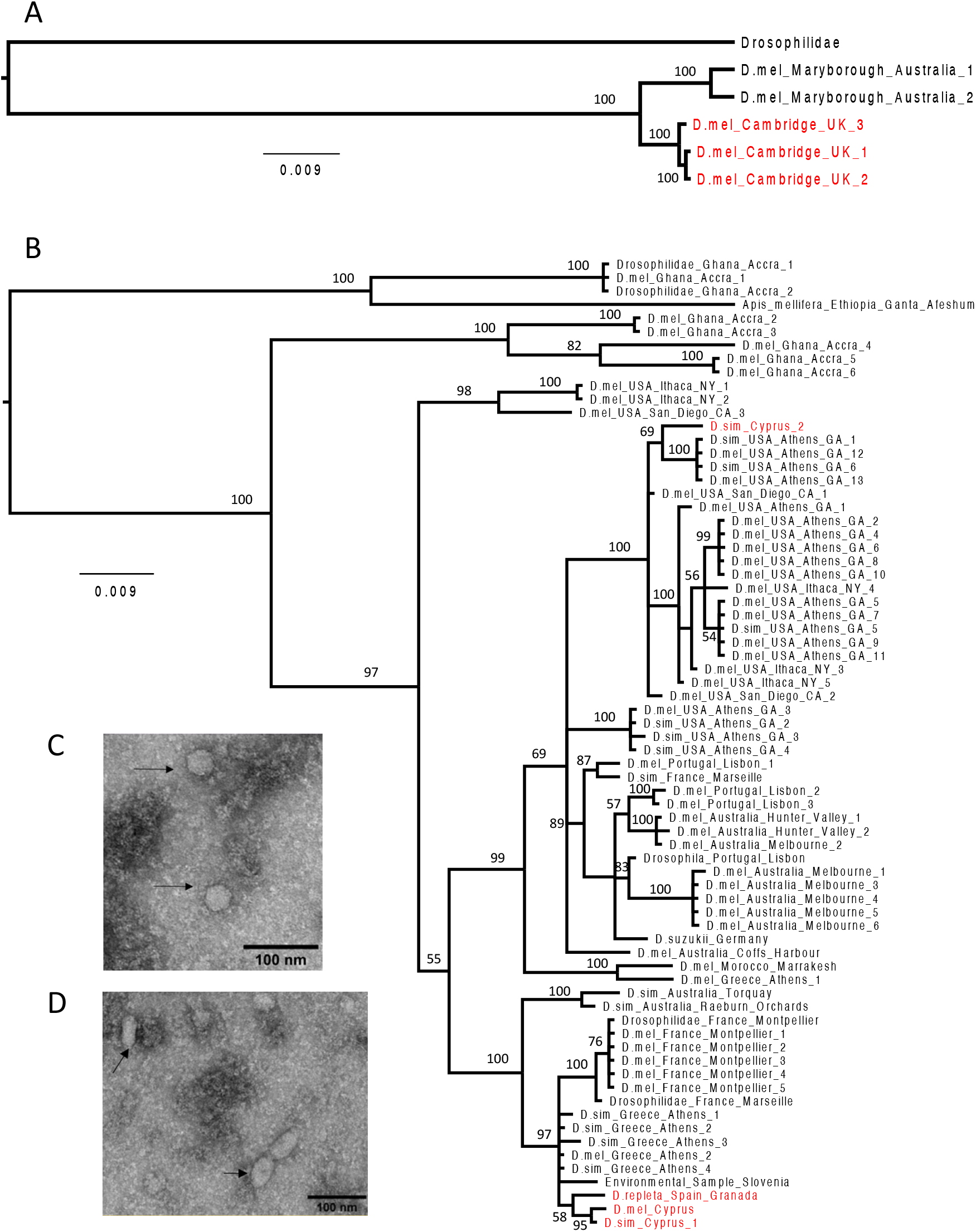
Phylogeny and morphology of LJV and NFV. Phylogeny was reconstructed using (A) whole-genome sequences of NFV and (B) partial sequences of the polyprotein gene of LJV. Nodes are labelled with Bayesian posterior probabilities (%) and the tree is midpoint rooted. Virus isolates from this study are red, and taxa are labelled by host species (Dmel, *D. melanogaster*; Dsim, *D. simulans*) and collection location. Accession numbers are in Table S2. Electron microscopy imaging of cells infected with (C) LJV isolate GBM-15052019-305 and (D) NFV isolate GBM-09102019-393 after negative staining. Arrows mark structures seen only in infected cells.

We used electron microscopy to examine the morphology of LJV and NFV virions. LJV showed an icosahedral viral particle (Figure 2C), consistent with other positive-sense single-stranded RNA viruses in the *Iflaviridae* family and previous reports of LJV [25]. Surprisingly, NFV repeatably showed elongated and spherical-shaped structures, which is consistent with this virus having pleomorphic virions (Figure 2D). Other viruses in the *Permutotetraviridae* have symmetrical icosahedral virions [26], so further work is required to confirm these are virions.

### Species-specific variation in susceptibility to NFV and LJV

To investigate the host range of LJV and NFV, we infected five species of *Drosophila* and measured viral titers after four days. The viral titers varied considerably depending on the combination of host and virus (Figure 3A; ANOVA, species × virus interaction: *F*=50.0, d.f.=4,40, *p*<10^−14^). For example, mean titres of LJV were more than 400 times greater in *D. simulans* than *D. sechellia*, despite these species being very closely related (Figure 3A). However, this pattern was reversed for NFV, where mean titres were over 30 times greater in *D. sechellia* than *D. simulans* (Figure 3A). There were similarly large differences among the other three species.

**Figure 3.**
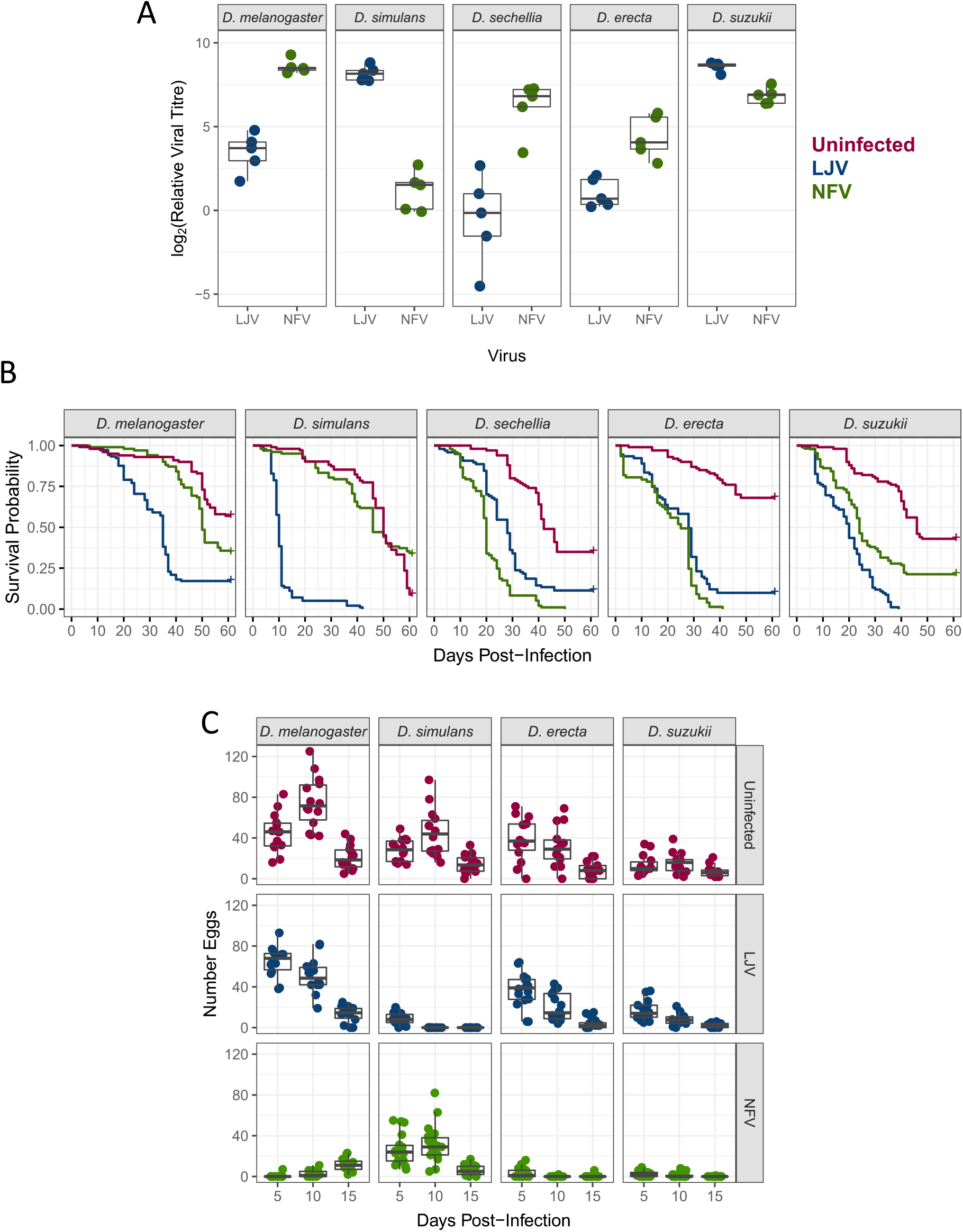
Host range and virulence of LJV and NFV. (A) Viral titre 4 dpi. Each point is a pool of 10-12 flies, and viral RNA levels are measured relative to *RpL32*. (B) Survival of flies after viral infection. Each line is from a mean of 98 flies. (C) Egg production after infection. Each point is the number of eggs laid of two females over 24h.

### NFV and LJV reduce lifespan

To evaluate the effect of viral infection on lifespan, we followed the survival of the five species of *Drosophila* after infection. With the exception of NFV in *D. simulans*, in all cases both NFV and LJV led to significantly increased mortality compared to uninfected flies (Figure 3B; Cox’s proportional hazards mixed effects model with Tukey’s adjustment: *p*<0.02 in all cases). However, the magnitude of this effect depended on the specific combination of host and virus. In *D. erecta* and *D. sechellia*, NFV caused significantly greater mortality than LJV (Figure 3B; Cox’s model with Tukey’s adjustment: *p*<0.002 in both cases). However, in the other three species this pattern was reversed, with LJV causing flies to die fastest (Figure 3B; Cox’s model with Tukey’s adjustment: *p*<0.001). In some species infection proved very virulent—for example, LJV had killed most *D. simulans* by 10 dpi (Figure 3B). In others, such as NFV infecting *D. melanogaster*, the virus caused only a modest increase in mortality.

There is an imperfect association between mortality rates and viral loads (comparison of Figures 3A and 3B). For example, *D. simulans* has very low titres of NFV and this virus did not reduce the survival of this species. However, this virus had higher titres in *D. melanogaster* than *D. erecta*, and yet caused greater mortality in the latter species.

### NFV can sterilize female flies

NFV infection caused strong reductions in female fecundity (Figure 3C). Combining data across the three timepoints we investigated, NFV caused a reduction in the number of eggs laid of 87% in *D. melanogaster*, 94% in *D. erecta*, 83% in *D. suzukii* and 41% in *D. simulans*. With the exception of the 5 dpi timepoint in *D. simulans*, these reductions were all statistically significant (GLM, contrasts with Tukey’s correction: *p*<=0.05 at every timepoint in each species). It is notable that this effect was markedly smaller in *D. simulans*, which is the species that had the lowest NFV titre (Figures 3A and 3C).

In three of the species, LJV caused considerably smaller reductions in fecundity than NFV— summing across timepoints, infection caused *D. melanogaster* to lay 8% fewer eggs, *D. erecta* 11% fewer, and *D. suzukii* 21% fewer (Figure 3C). The exception to this pattern was *D. simulans*, which suffered a 91% reduction in fecundity, and was completely sterile at 10 and 15 dpi (Figure 3C; GLM, contrasts with Tukey’s correction: *p*<0.001 at all timepoints). It is notable that LJV also causes very high mortality in *D. simulans* at the ages when we were measuring fecundity (Figure 3A).

### NFV infects the ovary and induces follicular degeneration

These results suggest that in some species NFV can have a specific impact on fecundity. For example, in *D. melanogaster* NFV causes less mortality than LJV, but causes a far greater reduction in fecundity. To investigate this further we examined the distribution of the two viruses in *D. melanogaster* ovaries using the monoclonal antibody J2 to detect viral dsRNA. A high level of dsRNA signal was observed in the ovaries of female flies infected with LJV and NFV (Figure 4). Interestingly, while LJV infection appears limited to the external muscular layers (epithelial sheet) that cover the ovariole, NFV infects the follicular epithelium, evidenced by the dsRNA signal detected within numerous follicle cells (Figure 4). Moreover, degenerated ovarian follicles with high levels of dsRNA signal can be observed at later stages, suggesting that oogenesis is severely affected in NFV-infected flies. Taken together, these results suggest that the viruses infect different tissues of the fly ovary producing different adverse effects.

**Figure 4.**
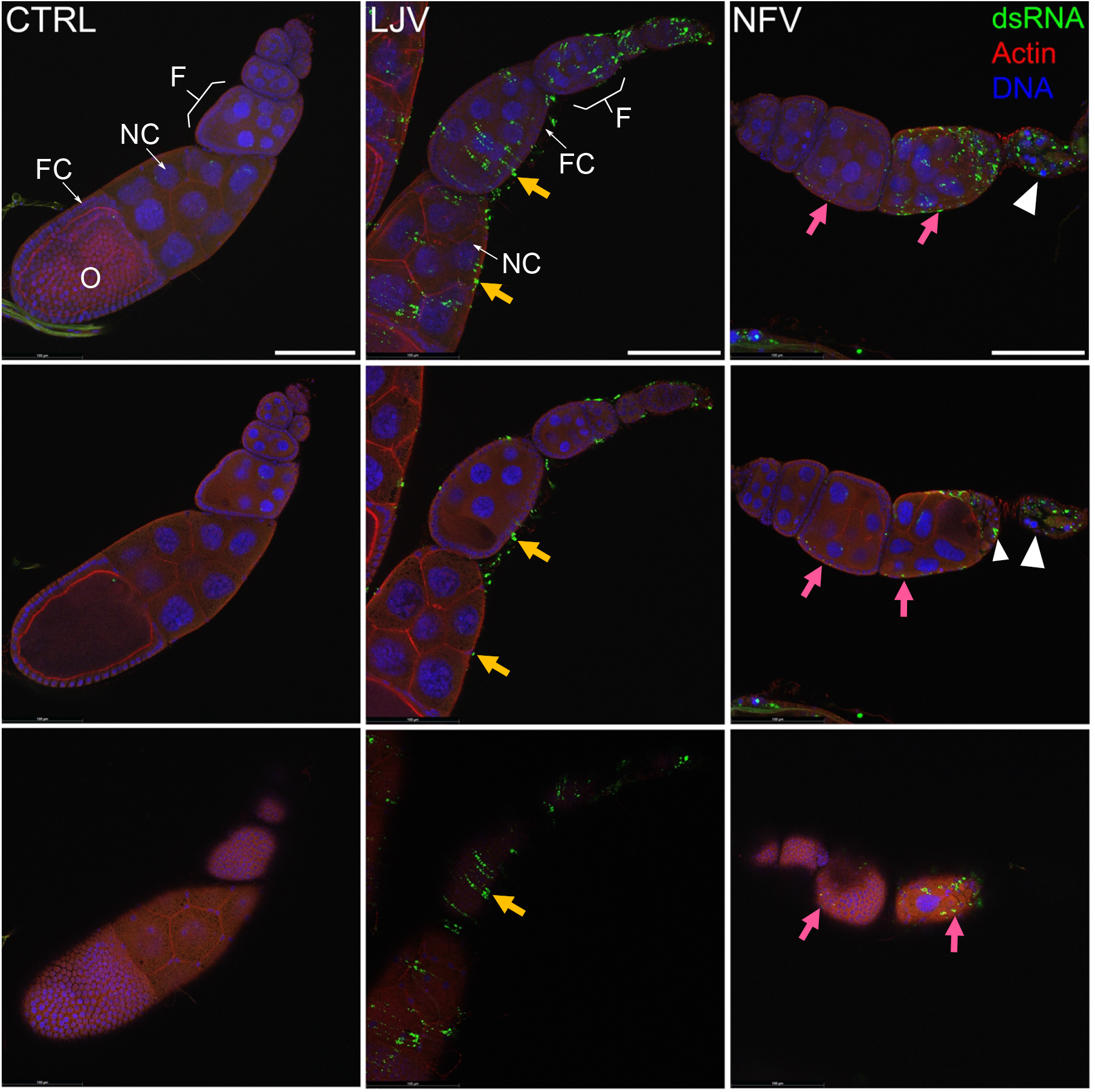
Immunostaining of ovaries showing the location of viral dsRNA. *D. melanogaster* ovaries were stained 10 dpi with DAPI to visualise DNA (blue), phalloidin to visualise the actin cytoskeleton (red), and mouse monoclonal antibody J2 to detect viral dsRNA (green), and imaged with a confocal microscope. From left to right, columns show individual ovarioles of females uninfected (CTRL), and infected with LJV and NFV. Top panels are a maximum intensity projection of 6 confocal planes taken from the same ovariole. Middle and bottom panels are individual confocal planes showing a transverse section of the ovariole (middle) or a superficial section (bottom). Note the dsRNA signal is absent from the CTRL, present at the ovariole surface (LJV, yellow arrows) or within the follicular epithelium (NFV, pink arrows). Letters represent oocyte (O), nurse cell (NC), follicle (F) and follicle cells (FC). In the case of NFV infection, infected follicles fail to progress through oogenesis and degenerate completely (white arrowheads). Scale bar indicates 100 µm.

### NFV is a potential biological control agent of D. suzukii

*Drosophila suzukii* is a major pest of commercially grown fruit, so the finding that NFV can partially sterilize this species means it is a potential biological control agent. We therefore tested whether these results held when flies were orally infected. Ten days after *D. suzukii* females were fed with yeast paste containing NFV, the virus could be detected in all the samples by qPCR (*N*=8, mean cycle threshold = 15). No virus was detected in the controls that had not been fed the virus (*N=*6, 40 PCR cycles). These results suggest that NFV could establish infections in populations of *D. suzukii* in crop fields.

We next tested whether NFV reduces the fecundity of *D. suzukii* after the flies have been infected orally. We found that NFV caused a 51% reduction in the number of eggs laid (GLM, main effect infection status: χ^2^=89, d.f.=1, *p*<10^−15^). This effect on fecundity was constant through time (GLM, infection status × time interaction: χ^2^=1, d.f.=3, *p*=0.79).

Finally, we investigated whether NFV-induced infertility could reduce the impact of *D. suzukii* on blueberry crops by exposing the fruit to infected and uninfected flies. As is the case in agriculture, we found that *D. suzukii* readily oviposit in blueberries and damage the fruit (Figure 5A). However, when blueberries were exposed to *D. suzukii* females that were orally infected with NFV, under half the number of flies emerged compared to fruit exposed to uninfected flies (Figure 5B; mean of 2.7 vs 6.95 flies/fruit; *t =* 5.18, *p* < 0.001). Taken together, the capacity of NFV to be orally acquired, and its subsequent effect on the number of adult flies emerging from fruit suggests it has the potential to be used as a biological control agent.

**Figure 5.**
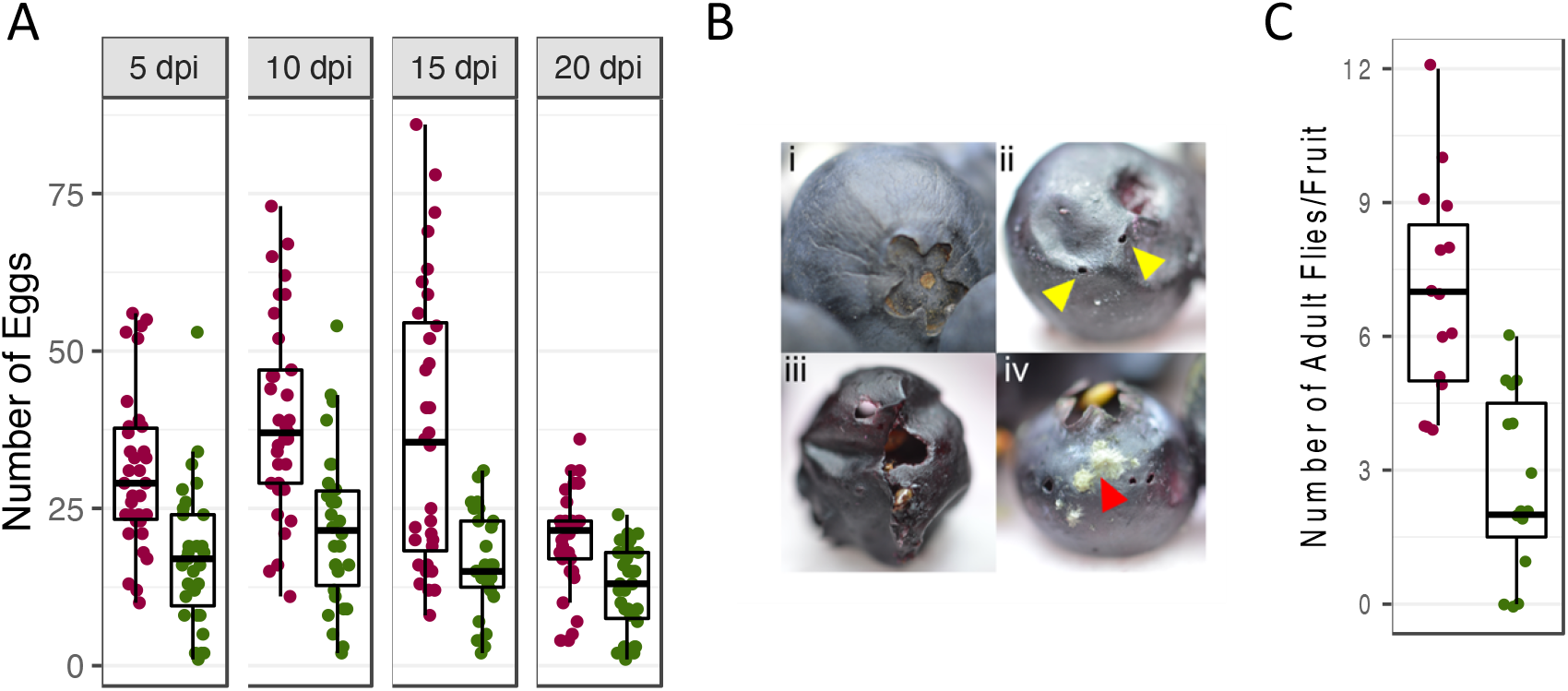
Effect of NFV on the fertility of the crop pest *D. suzukii*. (A) Fecundity of *D. suzukii* orally infected with NFV. Each point is the number of eggs laid by 5 females over 24h. (B) Blueberry fruit (i) before and (ii) after an infestation with *D. suzukii*, which uses a serrated ovipositor that pierces (yellow triangle) the skin to deposit eggs. The larvae consume the fruit from the inside (iii), and cause the proliferation of opportunistic microorganisms such as fungi (iv, red triangle). (C) The number of adult flies that hatch from blueberry fruits after being infected orally with NFV or fed a control solution. Each point is the number of flies that emerge from two fruits.

## Discussion

The discovery of an enormous diversity of insect viruses using metagenomics makes their isolation a pressing priority. We used a simple protocol to isolate viruses from *Drosophila*, which could easily be applied to any species where cell lines exist, and in any laboratory equipped for routine molecular- and micro-biology. We made a cell line that had greatly increased susceptibility to infection by suppressing anti-viral RNAi using the B2 protein. As B2 acts by binding viral RNA rather than a host factor [22], this should work in other insect species. After homogenizing wild-caught insects and adding them to these cells, we followed an approach that has been used in mosquitoes [21] and detected novel viruses from the presence of dsRNA, together with cytopathic effect in the infected cells. This allowed us to readily isolate LJV and NFV from multiple populations. In these experiments we selected only the cells with the strongest cytopathic effect and highest dsRNA concentration, so by relaxing these criteria in future studies we would anticipate that further viruses would be detected. Furthermore, some cells were coinfected with multiple viruses and this prevented us isolating KV. In future work this might be avoided if the wild flies are not pooled before adding them to the cell culture.

Both the viruses we isolated are positive-strand RNA viruses and belong to families of viruses that are widespread in insects. Using metagenomic sequencing and PCR screening, several studies have found that LJV is common in many wild *Drosophila* populations [5, 7, 11, 27–29]. It is a member of the *Iflaviridae* family, all members of which infect arthropods, including mosquitoes [30, 31], leafhoppers [32] and hymenopterans [33–35]. The pathology of other members of this family can range from harmful to asymptomatic [36]. Interestingly, LJV has been found in the Ethiopian honey bee (*Apis mellifera simensis*) [37], Australian honey bee (*A. mellifera*) [38], and the Asian hornet (*Vespa velutina*) [39], suggesting a relationship with hymenopteran viruses.

NFV is prevalent in both laboratory fly stocks and *Drosophila* cell cultures, but less common in wild populations [5]. Only two metagenomic studies have reported NFV in *Drosophila* wild populations. It belongs to the *Permutotetraviridae* family, which includes viruses that infect Lepidoptera [40], mosquitoes [41, 42], Hymenoptera [35], beetles [43], leafhoppers [44] and thrips [5, 45]. Viral sequences close related to NFV have been detected in a mosquito cell culture and *Aedes albopictus* mosquito samples collected from Sarawak, Borneo [46, 47].

While viral infection is known to be extremely common in natural *Drosophila* populations, for many viruses it is not known if these infections cause significant disease. We found that both NFV and LJV were virulent pathogens, capable of reducing both the survival and reproductive success of infected flies. This agrees with previous work on LJV in *D. suzukii*, which found a substantial reduction in the lifespan of infected flies. Given that LJV is estimated to infect ~9% of wild *D. melanogaster*, it is likely an important determinant of fitness [5].

There was a great variation in the effect of NFV and LJV viruses between species. Both viruses reduced survival. Female fertility was dramatically affected by NFV in all species except *D. simulans*. Conversely, LJV strongly reduced the fertility of *D. simulans* but had modest effects in other species. The mechanisms that explain these species-specific outcomes of host-virus interactions are unknown. However, the coevolution of hosts and pathogens can increase the natural variation of resistance, and drive differences between populations and species [48–50]. For example, the host range of the Nora virus is constrained by specific interactions between the host antiviral RNAi machinary and the viral suppressors of RNA silencing [51].

With the exception of *D. simulans* where titers are low, NFV had a strong sterilizing effect on females. In *D. melanogaster* this was associated with its ability to infect ovaries and cause ovarian follicles to degenerate. Many parasites have specific effects on host fertility, a trait often called parasitic castration [52]. This is sometimes interpreted as an adaptation of the parasite. If a parasite kills it host, it will die too. However, if it castrates the host, this may redirect resources to the parasite without reducing the period that the host is infectious [52]. Whether the sterilizing effect of NFV is an incidental byproduct of infection or an adaptation is unknown.

In surveys of natural *Drosophila* populations, it is common to find that virus infect multiple species but their prevalence varies [5], and this could be caused by ecology or differences in host susceptibility is unknown. We found that the viral load varied extensively, even among closely related species. The same was true for virulence, with the impact on fecundity and survival differing greatly among host species. More extensive surveys of DCV and sigma virus have revealed similarly large differences in the susceptibility of *Drosophila* species to infection [53–55]. Together, these results suggest that host genetics may be a key factor shaping the community of viruses in different *Drosophila* species.

NFV reduced the fecundity of *D. suzukii*, which is an invasive pest of fruit crops around the world. It was able to infect *D. suzukii* orally and reduce the number of flies that emerged from blueberry fruits, suggesting it merits investigation as a potential biological control agent. Other insect viruses have shown to cause mortality in *D. suzukii*, including Cricket Paralysis virus (CrPV), FHV, and DCV, DAV and LJV [6, 29, 56, 57] [25]. While these experiments suggest viruses might be suitable as biological control agents, more research is needed to determine whether producing and delivering RNA viruses is practical, or whether their effects on *D. suzukii* are sufficient to reduce crop damage.

Our results provides an alternative method to isolate *Drosophila* viruses from wild populations by suppressing *Drosophila* cells RNAi antiviral defence, detection of dsRNA, CPE inhibition assay and RNA sequencing. Additionally, we achieved the isolation of several isolates of LJV and NFV. We demonstrate that the virulence of these viruses varies greatly among closely related *Drosophila* species and depends on the specific combination between host species and virus. *Drosophila* is an important genetic and molecular model for the study of host-virus interactions, that together with these new viruses, may represent a useful tool for understanding the genetic and evolutionary basis of viral diseases.

## Material and Methods

### Fly collection

*Drosophila* flies were collected in the field at Limassol and Nicosia (Cyprus) in 2019, Budapest (Hungary) in 2018, Granada and Mallorca (Spain) during 2018–2019, Cambridge (United Kingdom) during 2019 (table 2.5.1). Flies were caught using an inverted cone fly trap containing banana baits sprinkled with dried yeast and ~300 µl of apple cider vinegar, or directly collected from decaying fruits using an entomological aspirator. Every 20–25 flies were transferred into a vial containing standard cornmeal diet and maintained at room temperature. Cyprus flies were transported to the Chrysoula Pitsouli’s laboratory at the University of Cyprus, where flies were anaesthetized using CO_2_ and split by species level under a stereomicroscope. *D. melanogaster* and *D. simulans* females are hard to identified morphologically, so that the females for these species were pooled together. All vials were transported to the Department of Genetics at the University of Cambridge and for further analysis.

### Virus isolation

Flies were anaesthetized using CO_2_, labelled, and sexed under the stereomicroscope. A pool of 5–10 females or males of the same species per sample were homogenized using a sterile pestle in an Eppendorf tube (1.5 ml) containing 200 µl of sterilized Ringer solution. Then, the tube was briefly spun and supernatant (~180 µl) was placed into a PVDF filter tube (Ultrafree-MC, 0.22 µm) and centrifuged at 1000g for 3 min to remove bacterial contamination and debris. The filtered solution was stored at −80°C for future analysis. Ten µl of the filtrate was inoculated into a well (96-well flat-bottomed plate) containing 190 µl of standard Schneider medium with DL2-B2 cells and incubated at 25°C. Three inoculations were performed as technical replicates per sample. The CPE of the cells was monitored daily for 10 days. On day 10, the plate was briefly spun (1000g for 30s) and 100 µl of the supernatant was stored at −80°C. The percentage of CPE per total well area (32.16 mm2) was scored from minimum (0) to maximum (3), where 0, 1, 2, and 3 represent 0-25%, 26-50%, 51-75%, and 75-100% of CPE, respectively. The remaining 100 µl was used in an enzyme-linked immunosorbent assay (ELISA) to detect double-stranded RNA (dsRNA). Samples with the highest CPE and dsRNA concentrations were retained.

### Enzyme-linked immunosorbent assay

One hundred µl of the cell culture was transferred into a 96-well microplate provided by the In-Cell ELISA kit (Abcam, ab111541), and treated following the manufacturer’s recommendations. For detecting dsRNA, the mouse monoclonal antibody J2 (Scicons) diluted 1:400 was used as primary antibody. As secondary antibody, goat anti-mouse IgG (IRDye® 800CW) diluted 1:1000 was used. The plate was stored at 4°C, and subsequently scanned on the Odyssey Infrared Imaging System using a fluorescent signal intensity, both 700 and 800 nm channels, and 4 mm focus. The fluorescence intensity of the image was analyzed on LI-COR Image Studio 4.0 Software for Odyssey. The microplate was normalized using the Janus green staining protocol as recommended by manufacturer instructions. Briefly, the microplate was emptied and 50 µl of 1 × Janus Green stain was added to every well for 5 min at room temperature, then the plate was washed 5 times with deionized water. Finally, 200 µl of 0.5 M hydrochloric acid was added to every well and incubated for 10 min. The microplate OD at 595 nm was measured in a microplate spectrophotometer SpectraMax iD3 (Molecular Devices).

### RNA extraction and quantitative polymerase reaction

Two ml of DL2-B2 cell culture (100,000 cell/ml) was grown in plastic culture flasks (see below, cell culture maintenance), and inoculated with 10 µl of the filtered solution from a sample positive for virus. After 8 days, the cell culture was transferred into a tube (15 ml) and briefly spun. Two hundred-fifty µl of the supernatant was used for total RNA extraction using the chloroform isopropanol TRIzol method following the manufacturer’s protocol (Life Technologies), the remaining supernatant was placed in cryogenic vials (Cryovial^®^) and stored at −80°C. One µl of RNA per sample was reverse-transcribed with Promega GoScript reverse transcriptase, followed of random hexamer primers, and then diluted 1:10 with nuclease free water. The complementary DNA (cDNA) was screened for the presence of the 14 *Drosophila* viruses: LJV, KV, DCV, Gown virus (GV), Thika 3 virus (TV), Galbut virus (GV), Drosophila-associated Partitiviridae (DPV), Nora virus (NV), Motts Mill virus (MV), Craigies Hill virus (CHV), Drosophila melanogaster Sigma virus (DMelSV), DAV, NFV and Chaq virus (CV). Additionally, the samples were screened for FHV and CrPv, which are occasionally found in fly stocks as contamination. For the qPCR, 1 µl of cDNA per sample was used to quantify the viral load by amplifying conserved regions of each virus genome (see primers, Table 2.5.2.), and the fly gene *RPL32* was used to normalize the expression. qPCR was performed on an Applied Biosystems StepOnePlus system using Sensifast Hi-Rox Sybr kit (Bioline) with the following PCR cycle: 95°C for 2min followed by 40 cycles of: 95°C for 5 secs followed by 60°C for 30 secs. Two technical replicates per reaction were carried out per sample with both the target virus and reference genes. To estimate the relative viral load, the formula ΔΔCt = ΔCt_*rpl32*_ –ΔCt_target *virus*_ was used, where ΔCt_*rpl32*_ and ΔCt_target *virus*_ represent the mean of the technical replicates.

### Purification of viral particles

To purify the viral particles, we used the end-point-dilution. The stock samples with virus detected were thawed on ice, and 10 µl of it inoculated into a well (96-well flat-bottomed plate) containing medium and DL2-B2 cells. Eight replicates of each sample were 1:10 serially diluted 12 times, and the CPE was monitored daily for 10 days. This process was repeated twice. The serially diluted samples were then grown in 6 ml of cell culture (6-well flat-bottomed plate) for 5 days.

### Library preparation

One Illumina sequencing library per sample was prepared, including one library for the virus-free DL2-B2 cells used in the isolation protocol as control. One µg of total RNA was used for the NEBNext® Ultra™ Directional RNA Library Prep Kit (New England Biolabs) without removing ribosomal RNA. cDNA synthesis was performed according to NEB protocol, and NEBNext Adaptor with hairpin loop structure were ligated to prepare for hybridization. Size selection was performed using NEBNext Sample Purification Beads (New England Biolabs), and indexes were added using NEBNext Multiplex Oligos for Illumina (New England Biolabs). The library quality was evaluated with the Agilent Bioanalyzer system. Libraries were transported to the Cancer Research UK Genomics Core Facility and sequenced on NovaSeq 6000 system (Illumina).

### Bioinformatics

The raw data quality assessment and adapter removal were performed with TrimGalore. Then, sequencing reads were mapped to both the *D. melanogaster* genome and all published *Drosophila* associated-virus sequences available in the National Center for Biotechnology Information (NCBI) [47] using STAR [48]. Subsequently, the unmapped reads were mapped to the SILVA rRNA database [49] to remove all rRNA using bowtie2 [50]. *De novo* assembly of the unmapped reads was performed with TRINITY software [59], together with the utility TransDecoder to identify coding regions within the transcripts with ORFs higher than 30 amino acids. These contigs were then queried against NCBI RNA virus proteins using blastx to identify putative viruses. The virus-related contigs were confirmed by querying the non-redundant protein database RefSeq by blastx.

To obtain the genome sequence of viruses that had been isolated we mapped reads to published genomes as described above and genetic variants were called with the program Freebayes. These variants were then used to modify the published genome sequence.

### Phylogenetic analysis

All sequences of NFV and LJV were downloaded from NCBI for phylogenetic analysis. Complete genome sequences of NFV (~4761 bp) and LJV partial length sequences (~690 bp) of the polyprotein gene were aligned with MUSCLE [60] using default settings. The best-fit nucleotide substitution model for the aligned sequences was determined by jModeltest [61]. Bayesian inference phylogenetic analysis was performed in MrBayes version 3.2.2 [62], with a General Time Reversible (GTR) DNA substitution model (lset nst= 6) Gamma distributed (rates=gamma), 4 independent chains (nchains =4) and recording the tree and parameters every 1000 generations (samplefreq= 1000). Several runs were performed to reach the optimal range of convergence adjusting the temperature parameter for heating the chains (temp = 0.1). All analyses consisted of 10 million generations. Sixty-four sequences were added to the analysis from the GenBank database from previous studies for comparison.

### Transmission electron microscopy

Virus samples with LJV and NFV were inoculated in DL2-B2 cell culture for 4 days. Two hundred µl of the culture was placed into a PVDF filter tube (Ultrafree-MC, 0.22 µm) and centrifuged at 1000g for 3 min. The supernatant was centrifuged twice using different filter tubes to eliminate the cell debris. Aliquots of the supernatant samples were transported to the Cambridge Advanced Imaging Centre for imaging processing. Samples were adsorbed onto glow-discharged 400 mesh copper/carbon film grids (EM Resolutions) for about 1 min. Then, transmission electron microscopy (TEM) grids were passed over two drops of deionized water to remove any buffer salts and stained in 2% (w/v) aqueous uranyl acetate for about 30 seconds. Uranyl acetate dye was drained off the TEM grid using filter paper and grids were allowed to air dry. Samples were viewed using a Tecnai G20 transmission electron microscope (FEI/ThermoFisher Scientific) run at an accelerating voltage of 200 keV using a 20-µm objective aperture to improve contrast.

### Cell culture maintenance

*Drosophila* cultured cell lines DL2 and DL2-B2 were used in the experiments. Cells were grown in plastic culture flasks containing Schneider medium supplemented with 10% heat-inactivated fetal bovine serum (FBS) and streptomycin 100 µg/ml and penicillin 100 U/ml to inhibit bacteria and fungus contamination. Culture flasks were passaged every 6 days and incubated at 25 °C.

### Amplification and digestion of FHV B2 open reading frame

FHV cDNA was synthesized from RNA isolated from infected flies using GoScript Reverse Transcriptase system (Promega), the resulting cDNA was diluted 2x before use in a PCR reaction. The open reading of the FHV B2 viral suppressor of RNAi [63] was amplified from FHV cDNA using FHV-B2-not_Fw (5’GCACGCGGCCGCACCATGCCAAGCAAACTCGCGCTAATC’3) and FHV_B2_not_Rv (3’GCACGCGGCCGCCCCAGTTTTGCGGGTGGGGGGTC’5) primers. FHV_B2_not_Fw incorporated a NotI restriction site and consensus Kozak sequence at the 5’ end of the open reading frame. FHV_B2_not_Rv incorporated a NotI restriction site, 2 extra bases to put the open reading frame in-frame with the V5 and His epitope tags in the vector and removed the stop codon at the 3’ end. Q5 hot start DNA polymerase (New England Biolabs) was used in the amplification reaction with a touchdown thermocycling protocol. A 5µl aliquot of the resulting 345 bp PCR product was analyzed by agarose gel electrophoresis. The remainder was purified using the QIAquick PCR Purification Kit and the B2 amplicon was eluted in 90µl of 1 mM Tris-HCl pH 8.0. The fragment was then digested by adding 10µl of 10x sure cut restriction enzyme buffer and 1.5µl (15 units) of NotI restriction enzyme (New England Biolabs) and incubating at 37°C for 1:45 h. The digested fragment was purified by gel electrophoresis and the QIAquick gel extraction kit (Qiagen) being careful to not expose the DNA to the fluorescent light for more time than was necessary and eluted in 20 µl of 1mM Tris-HCl pH 8.0.

### Plasmid digestion

Meanwhile, 150 ng of pMT-puro plasmid was digested with NotI restriction enzyme in a total reaction volume of 100 µl at 37°C for 1:45 h. To prevent re-annealing of the empty plasmid during ligation the 5’ ends of the digested plasmid were dephosphorylated by adding 1ul of alkaline phosphatase and incubation at 37°C for 1 h. The digested plasmid was then purified using a PCR purification kit (Qiagen) according to the manufacturer’s recommendations and eluted in 20µl of 1 mM Tris-HCl pH 8.0.

### Ligation and transformation

Plasmid and B2 insert were ligated together using a rapid ligation kit (Promega). Ligation reactions contained 2.5µl 4x buffer, 0.5µl T4 ligase, 1µl vector, 1µl insert and were incubated at room temperature for 15 min. A control ligation that contained plasmid only was also set up. The ligations were transformed into 50 µl of chemically competent sub-cloning grade *E. coli* DH5a strain (New England Biolabs). Cells were thawed on ice then 2µl of the ligations were added to the cells, which were then carefully mixed using gentle agitation and returned immediately to ice and incubated for 30 min. The cells were then heat shocked for 30 s at 43°C in a water bath and momentarily returned to ice. Nine hundred and fifty µl of Luria-Bertani (LB) was gently added to the cells and they were incubated with shaking at 37°C for 45 min. The cells were pelleted by centrifugation at 6000 g for 1min and all but 200 µl of the supernatant was removed. The cells were re-suspended in the remaining LB, and they were spread onto 96 mm diameter bacterial LB agar plates containing 10 ug/ml ampicillin and the plates were incubated inverted at 37°C overnight.

### Analysis of clones

Next day 30 single colonies were removed using a pipette and washed off into 10 µl LB containing 10 ug/ml ampicillin. These cultures were tested for the presence and orientation of the inserts by PCR. For each culture 3 PCR reactions were set up in a 96-well PCR plate, primer pairs were as follows: FHV_B2_not_Fw/FHV_B2_not_Rv; pMT_forward/FHV_B2_not_Rv; pMT_forward/FHV_B2_not_Fw. Thermopol DNA polymerase was used (New England Biolabs) and a touchdown PCR protocol. 0.3 µl of each culture was used as a template in a 15 µl reaction volume. Amplified fragments were analyzed by gel electrophoresis. The remainder (~9 ul) of the 5 cultures containing plasmids with B2 inserts of the correct orientation were used to inoculate 10 ml LB cultures and were incubated overnight with shaking at 37°C. The following day plasmids were purified from these cultures (QIAprep Spin Miniprep Kit, Cat No./ID: 27104) and quantified using Qubit hsDNA assay. Two hundred ng of each plasmid were digested with 0.5 µl of NotI in a 10 µl reaction volume for 1 h at 37 °C and then analyzed by gel electrophoresis. All plasmids contained the correct sized insert. Three of the 5 plasmids were sequenced using pMT_forward and BGH_reverse primers. Each sequencing reaction contained 200 ng of template plasmid DNA. All 3 plasmids had the correct sequence.

### DL2 cell transfection

One day grown DL2 cells were transfected using the plasmids containing the B2 insert. Transfection was performed using Effectene transfection (QIAGEN) reagent according to the manufacturer’s instructions. Transfected cells were incubated in 1.6 ml Schneider medium containing ~1.8×10^6^ DL2 cells seeded in 6-well plastic plates. Growth medium was supplemented with antibiotics and incubated as mentioned above. Second-day post-transfection, the medium was removed and replaced with medium containing 10 µg/ml of puromycin and incubated for 3 days. Each well was monitored every 24 h to check cell conditions. To induce the expression of the B2 protein, 5 mM of CuSO4 was added 24 h before the experiments.

### Immunohistochemistry

Immunohistochemistry was performed to test the expression of the vector. pMT contain the V5 epitope tag allowing its rapid detection with Anti-V5 Antibody. DL2-B2 induced and uninduced cells (CuSO4-free) were cultured under the conditions mentioned before. Briefly, 1 ml of medium was centrifuged at 2000g for 3 min, the supernatant medium was discarded and washed twice with 1% v/v phosphate-buffered saline (PBS). Then, 20 µl of the cells were placed on a slide coated with Poly-L-Lysine for 30 min. Every well was washed 3x with 1% v/v PBS, and 20 µl of 4% w/v paraformaldehyde was added for 30 min, followed by blocking and permeabilization with 1× PBS, 0.01% v/v Triton-X (Sigma), 1% v/v Normal Goat Serum (NGS) for 30 minutes. The cells were labelled with V5 tag mouse monoclonal antibody (Thermofisher) at a dilution of 1:1000 in 1% v/v PBS and incubated overnight at 4°C, then labelled with Alexa Fluor® 488 rabbit anti-mouse IgG secondary antibody (Thermofisher) at a dilution of 1:1000 and incubated overnight at 4°C. Finally, cells were washed 3x with 1% PBS and stained with Hoechst for 5 min and washed 3x with PBS. Mount slides with 80% glycerol and seal edges with nail polisher. The slide was mounted with 80% glycerol and edges were sealed with nail polisher. The images were taken on a Nikon Eclipse 90i microscope at 20× magnification and processed using the Fiji software.

### Cell count and dsRNA after viral infection

To test the susceptibility of the DL2-B2 cells, we counted the number of viable cells after infection with DCV-C. DL2-B2 or DL2 cells (100,000 cell/ml) were grown in a 96-well flat-bottomed microplate. Each well contained 90 µl of Schneider medium, puromycin 10 µg/ml and CuSO4 5 mM. After 24 h, wells were randomly inoculated with 10 µl of DCV at different concentration treatments. All protocols and cell culture incubation were performed at 25°C. Cell concentration was estimated 72-, 144-, and 216-hours post-infection (hpi) by pipetting ~20 µl of cell solution into a disposable hemocytometer (FastRead 102) under the microscope.

To quantify dsRNA in DL2-B2 and DL2 cells, cells were infected with DCV or FHV and 48 hpi, cells were transferred into a 96-well microplate provided by the In-Cell ELISA kit, and the dsRNA was detected and analyzed as mentioned above.

### Drosophila species and infection of flies

Five species that belong to the subgenus Sophopora group were used in the study. Flies were maintained in glass vials (~ 28.5 × 95 mm) with standard cornmeal (1200 ml water, 13 g agar, 105 g dextrose, 105 g maize, 23 g yeast, 35 ml Nipagin 10% w/v), and incubated at 25°C and ~70% humidity.

The infection experiments used LJV isolate GBM-15052019-4-305 that was isolated from Gialousa, Cyprus, and NFV isolate GBM-09102019-1-393 that was isolated from Cambridge, UK. All virus isolates were cultured in DL2-B2 cells. To produce aliquot of virus for the experiments, 2 ml of cell culture was added to a conical tube (Falcon™) and briefly spun (1000g for 2 min) to remove cell’s debris, then the supernatant was aliquoted (10 µl) into sterilized 0.2 ml PCR tubes and stored at −80°C. The viral concentration of the isolates was estimated using the TCID50 ml^−1^ method [64]. To do this, DL2-B2 cells (100,000 cell/ml) were growth in a 96-well flat-bottomed plate containing 190 µl of standard Schneider medium supplemented as mentioned above. Then, 10 µl of the viral solution was inoculated into a well, and serially diluted 1:10 14-fold. The cytopathic effect of the cells was monitored daily for 10 days. The aliquot stocks were thawed immediately before each pricking assay on ice and diluted in sterile Ringer’s solution to standardise the concentration of the virus isolates to 1×10^5^ TCID50 ml^−1^.

### Viral infection

Flies were anesthetized on a CO2 pad and then pricked on the dorsolateral thorax under a stereomicroscope using a needle (Austerlitz Insect Pin) dipped into a Ringer’s solution containing 1×10^5^ TCID50 ml^−1^ viral titre. To avoid cross-contamination between viruses, different CO2 pads and needles were used for each virus. These utensils were kept in independent plastic bags and cleaned with Virkon® (5% w/v) and ethanol (70% v/v) frequently. After infection, infected flies were kept in independent trays for each virus treatment to avoid cross-contamination. Unless otherwise mentioned, flies were transferred every 3 days to new vials with fresh food cornmeal and incubated at 25°C over the course of the experiments.

### Oral infection

To infect orally the flies, a cohort of virgin adult females 3–5 days old was transferred into an empty vial without food containing a damp towel paper for 24 h at 25°C. The next day, the starved females were transferred into a vial containing 300 µl of yeast paste (25% w/v yeast powder, 5% vegetable red dye v/v, and 1×10^5^ TCID50 final concentration of one of the viruses) and incubated at 25°C. Another cohort of females was transferred into vials with Ringer’s solution (25% w/v yeast powder and 5% red dye v/v) as a control. To confirm that the flies ingested the virus solution, the next day flies’ gut was checked under a stereoscopic. Flies with red-stained intestines were selected for the experiment and transferred into new vials with fresh cornmeal food.

### Virus quantification

The total RNA of the homogenized flies was extracted using the chloroform isopropanol method following the manufacturer’s protocol (Life Technologies). One µl of RNA per sample was reverse-transcribed with Promega GoScript reverse transcriptase using random hexamer primers, and then diluted 1:10 with nuclease-free water. qRT–PCR was performed on an Applied Biosystems StepOnePlus system using Sensifast Hi-Rox Sybr kit (Bioline) with 1 µl of complementary DNA (cDNA) per sample was used to quantify the viral load using specific primers for LJV LaJolla1_foward (5’-CGGACCAGAGTGTAGCCAAG-3), and LaJolla1_reverse (5’-AGTGCCATCCAYCGATTTGT-3’), and NFV NewfiledVirus_2_forward (5’-TTGATGATGTCGCCACGAGA-3’), NewfiledVirus_2_reverse (5’-CATTCGCCGAGACCTCCATC-3’). The fly gene *RPL32* was used to normalize the expression using primers RpL32_forward (5’-TGCTAAGCTGTCGCACAAATGG-3’) and RpL32_reverse (5’-TGCGCTTGTTCGATCCGTAAC-3’) [65]. The qRT-qPCR was performed with the following PCR cycle: 95°C for 2min followed by 40 cycles of 95°C for 5 secs followed by 60°C for 30 secs. Two technical replicates per qRT-PCR reaction were carried out per sample with both the viral and reference genes. To estimate the relative viral load, the formula ΔCt = Ct_*RPL32*_–Ct_target_virus_ was used, where Ct is the mean Ct value of the technical replicates performed on each target sequence.

### Viral titre across different Drosophila species

To evaluate the effects of the viruses across different fruit flies, 5 *Drosophila* species were infected with LJV and NFV. For each species, two male and 2 female flies were transferred into 6 vials (n = 12 vials total), containing standard diet and incubated at 25°C— flies were discarded at the 2^do^ days. After 2 weeks, 10 adult males (3–5 days old) for each species were transferred into a vial with fresh cornmeal food. Then, 6 vials per species treatment were pricked with either LJV or NFV. Three vials were pricked with Ringer’s solution as an uninfected control. Four-days dpi, all flies per vial were anaesthetized using CO2 and transferred into Eppendorf tube (2 ml) containing beads. Every tube was chilled on ice for 10–15 min and 250 µl of TRIzol® Reagent (Invitrogen) was added. Immediately, tubes were homogenised using a Qiagen TissueLyser II and stored at −80°C. For each tube, the total RNA was extracted, and the relative virus concentration was estimated as mentioned above.

### Lifespan

To assess the effect of the viruses on lifespan of flies, different *Drosophila* species were infected with either LJV or NFV. For each species, 12 vials with flies (2 males and 2 females) were set up and the parental flies discarded as in the experiment above. After 2 weeks, 20–25 adult males (3–5 days old) per species were transferred into a vial with fresh cornmeal food. Four vials per species were pricked with either LJV, NFV or Ringer’s solution (control). The number of dead flies per vial was monitored daily until the last fly was dead. Mortality on day 1 was attributed to the damage induced by the needle, and the data was discarded from the analysis.

### Fecundity

To investigate the effect the viruses on fecundity after viral infection, different female fly species were infected with LJV and NFV. For each species, vials with flies (2 males and 2 females) were set up and the parents were discarded as mentioned above. One virgin female and 2 males (3–5 days old) per species were transferred into a vial with cornmeal food with live yeast sprinkled. Females were pricked with either NFV, LJV, or Ringer’s solution (control). Males were not infected. To quantify the number of eggs produced, two females and 2 males were transferred into new vials without yeast with fresh food and allowed to lay eggs for 24h. The number of eggs was quantified under a stereomicroscope at 3-time points, between 4–5, 9–10, and 14–15 dpi. Because *D. sechellia* fertility is reduced in standard *Drosophila* diet, it was not used in this experiment.

### NFV viral oral infection of D. suzukii

To evaluate the fertility of *D. suzukii* females after oral viral infection over time, *D. suzukii* female flies were infected with NFV. This experiment was performed as the abovementioned fecundity assay with the exception that females were infected orally. Virgin females (3-5 days old) were orally infected with NFV or Ringer’s solution (control) and transferred into new vials with fresh food and 2 males 3–5 days old. The number of eggs produced was quantified at 4-time points, between 4–5, 9–10, 14–15, and 19–20 dpi.

To investigate whether NFV can establish an oral viral infection on *D. suzukii* flies, female flies were infected with either NFV or a control solution. Adult females 3–5 days old were orally infected with NJV or Ringer’s solution as the abovementioned oral infection protocol. Flies were transferred into new vials with fresh food and ten days after oral infection, the total RNA of single female flies was extracted, and the relative virus concentration was estimated as mentioned above.

NFV-induced fertility in *D. suzuki* was determined when allowed to oviposit on blueberry fruits. Female flies were orally inoculated with either NFV or a control solution, as mentioned above. The females were kept in vials with fresh cornmeal food for 4 days. Then, 5 infected females and 5 uninfected males were placed in a vial containing 2 organic blueberry fruits and a damp towel paper to maintain humidity. After 10 days, the flies were discarded and the number of flies that emerged from each vial was measured for 15 days.

### Statistical Analysis

We conducted all statistical analyses in R [66]. All graphs were generated with the ‘ggplot2’ package [67]. Experiments were analysed using an ANOVA had normality and homoscedasticity of the data was detected with the Shapiro Test and Levene Test respectively, and data was log or squared root transformed when assumptions were violated. *P* values from post-hoc comparisons were adjusted by the Tukey method using the ‘lsmeans’ package [68]. To analyse mortality, a Cox’s proportional hazard mixed-effect model was fitted in the ‘coxme*’* package [69], including the fixed effects of *Drosophia* species, virus treatment (LJV, NFV, or control) and their interaction, and vial of flies as a random effect.

Fecundity was analysed Generalized Linear Model with a negative binomial distribution, including zero inflation where appropriate. This was fitted using the ‘glmmTMB’ package [70], including the fixed effects of *Drosophia* species, virus, and timepoint as fixed effects (together with interactions). To analyse the number of flies hatched between blueberry fruits infected and uninfected with *D. susukii*, a *t* test was used. A Type III Wald test was used to estimate significance of effects within models using the ‘Anova()’ function in the ‘car’ package [71].

## Acknowledgements

Chrysoula Pitsouli kindly allowed us to use her laboratory in Cyprus. Ben Longdon and Luis Teixeira supplied fly stocks.

## Data Availability

Viral genome sequences have been submitted to GenBank (Accession Numbers: OP263078-OP263084). The RNAseq reads have been deposited in the NCBI Sequence Read Archive (bioproject PRJNA872469, accessions SAMN30466843-SAMN30466854). The raw data and scripts to reproduce figures and statistical analyses have be submitted to the Cambridge Data Repository (doi xxxxxxavailable on acceptance xxxxxxxx).

---

**Table S1.** Virus detected by qPCR in 19 samples showing high cytopathic effect and high dsRNA concentration.

| Sample ID | Place | Country | Host | N flies | Sex | Virus |
| --- | --- | --- | --- | --- | --- | --- |
| gb-08052019-2-26 | Gialousa | Cyprus | *D. simulans* | 5 | females | DCV |
| gb-09052019-3-59 | Plantation | Cyprus | *D. simulans* | 5 | females | LJV |
| gb-09052019-3-60 | Plantation | Cyprus | *D. simulans* | 5 | females | LJV |
| gb-08052019-5-91 | Gialousa | Cyprus | *D. simulans* | 5 | females | LJV |
| gb-08052019-5-93 | Gialousa | Cyprus | *D. simulans* | 5 | females |  |
| gb-08052019-5-98 | Gialousa | Cyprus | *D. simulans* | 5 | males | LJV |
| gb-10052019-2-112 | Gialousas | Cyprus | *D. repleta* | 5 | females |  |
| gb10052019-3-126 | Gialousas | Cyprus | *D. simulans* | 5 | females |  |
| gb10052019-3-127 | Gialousas | Cyprus | *D. simulans* | 5 | females |  |
| gb-11052019-3-130 | Gialousas | Cyprus | *D. immigrant* | 6 | females |  |
| gb-11052019-2-131 | Gialousas | Cyprus | *D. immigrant* | 5 | females | DCV |
| gb15052019-4-305 | Gialousa | Cyprus | *D. melanogaster* | 7 | males | LJV |
| gb15052019-4-306 | Gialousa | Cyprus | *D. simulans* | 7 | males | LJV |
| GOY-2105209-1-311 | Granada | Spain | *D. virilis* | 4 | females | LJV |
| gb-09102019-1-386 | Cambridge | UK | *D. melanogaster* | 10 | males | NFV, LJV |
| gb-09102019-1-390 | Cambridge | UK | *D. melanogaster* | 5 | males |  |
| gb-09102019-1-391 | Cambridge | UK | *D. melanogaster* | 5 | males | LJV |
| gb-09102019-2-392 | Cambridge | UK | *D. melanogaster* | 5 | females | NFV, Kallithea |
| gb-09102019-1-393 | Cambridge | UK | *D. melanogaster* | 7 | females | NFV, Kallithea |

**Table S2.**
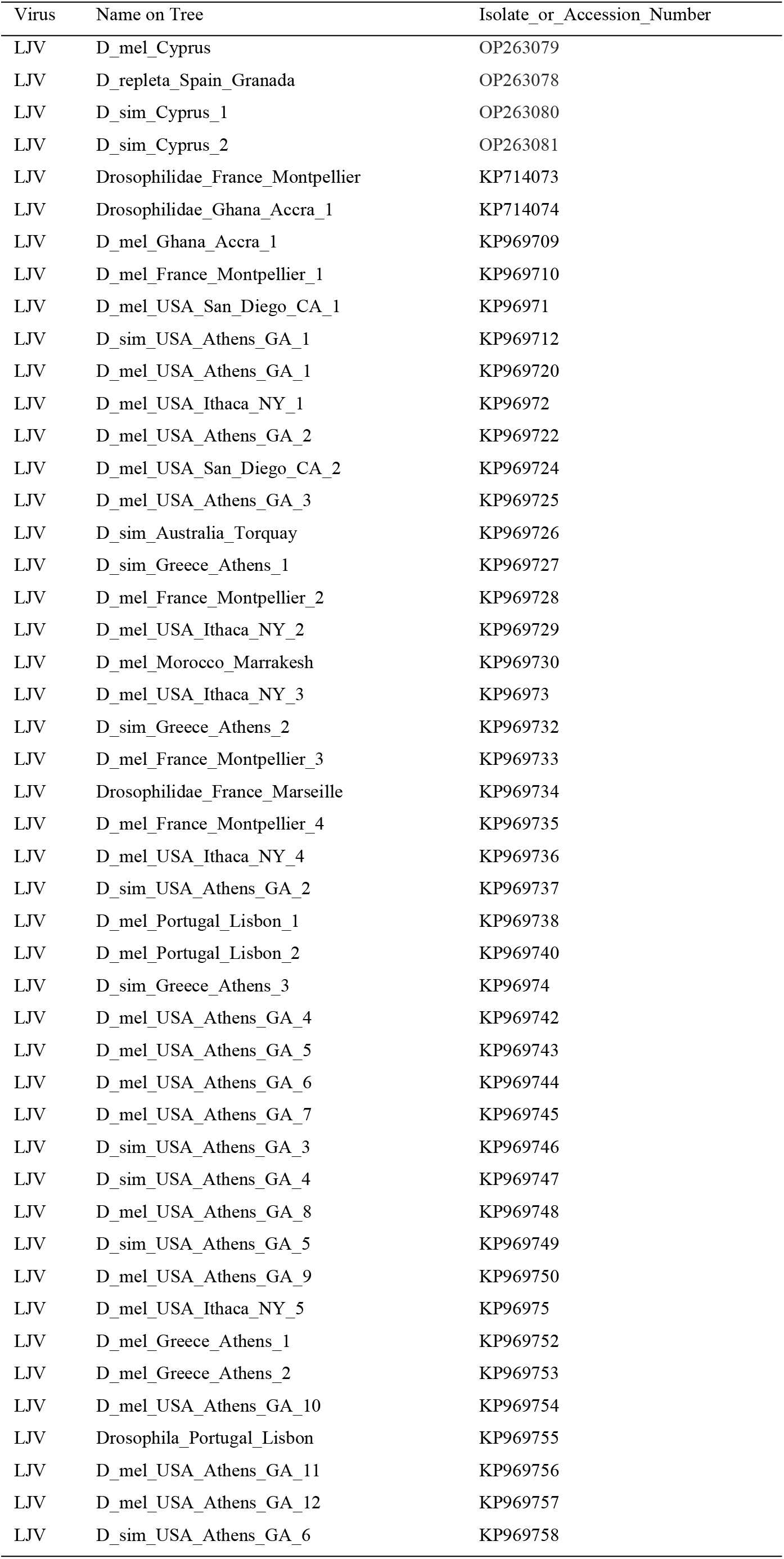
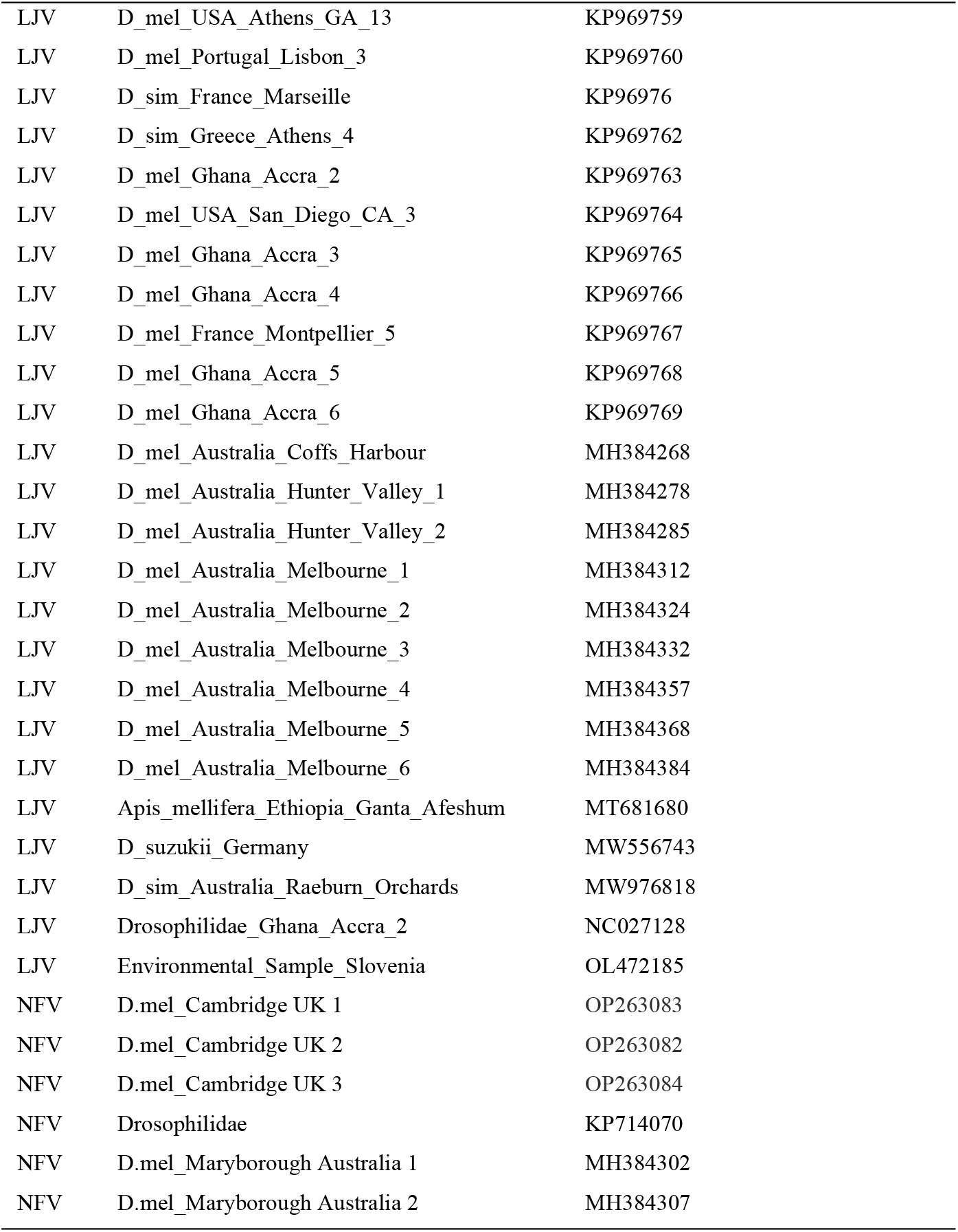
Accession and Isolate Numbers of Virus Sequences.

| Virus | Name on Tree | Isolate_or_Accession_Number |
| --- | --- | --- |
| LJV | D_mel_Cyprus | OP263079 |
| LJV | D_repleta_Spain_Granada | OP263078 |
| LJV | D_sim_Cyprus_1 | OP263080 |
| LJV | D_sim_Cyprus_2 | OP263081 |
| LJV | Drosophilidae_France_Montpellier | KP714073 |
| LJV | Drosophilidae_Ghana_Accra_1 | KP714074 |
| LJV | D_mel_Ghana_Accra_1 | KP969709 |
| LJV | D_mel_France_Montpellier_1 | KP969710 |
| LJV | D_mel_USA_San_Diego_CA_1 | KP96971 |
| LJV | D_sim_USA_Athens_GA_1 | KP969712 |
| LJV | D_mel_USA_Athens_GA_1 | KP969720 |
| LJV | D_mel_USA_Ithaca_NY_1 | KP96972 |
| LJV | D_mel_USA_Athens_GA_2 | KP969722 |
| LJV | D_mel_USA_San_Diego_CA_2 | KP969724 |
| LJV | D_mel_USA_Athens_GA_3 | KP969725 |
| LJV | D_sim_Australia_Torquay | KP969726 |
| LJV | D_sim_Greece_Athens_1 | KP969727 |
| LJV | D_mel_France_Montpellier_2 | KP969728 |
| LJV | D_mel_USA_Ithaca_NY_2 | KP969729 |
| LJV | D_mel_Morocco_Marrakesh | KP969730 |
| LJV | D_mel_USA_Ithaca_NY_3 | KP96973 |
| LJV | D_sim_Greece_Athens_2 | KP969732 |
| LJV | D_mel_France_Montpellier_3 | KP969733 |
| LJV | Drosophilidae_France_Marseille | KP969734 |
| LJV | D_mel_France_Montpellier_4 | KP969735 |
| LJV | D_mel_USA_Ithaca_NY_4 | KP969736 |
| LJV | D_sim_USA_Athens_GA_2 | KP969737 |
| LJV | D_mel_Portugal_Lisbon_1 | KP969738 |
| LJV | D_mel_Portugal_Lisbon_2 | KP969740 |
| LJV | D_sim_Greece_Athens_3 | KP96974 |
| LJV | D_mel_USA_Athens_GA_4 | KP969742 |
| LJV | D_mel_USA_Athens_GA_5 | KP969743 |
| LJV | D_mel_USA_Athens_GA_6 | KP969744 |
| LJV | D_mel_USA_Athens_GA_7 | KP969745 |
| LJV | D_sim_USA_Athens_GA_3 | KP969746 |
| LJV | D_sim_USA_Athens_GA_4 | KP969747 |
| LJV | D_mel_USA_Athens_GA_8 | KP969748 |
| LJV | D_sim_USA_Athens_GA_5 | KP969749 |
| LJV | D_mel_USA_Athens_GA_9 | KP969750 |
| LJV | D_mel_USA_Ithaca_NY_5 | KP96975 |
| LJV | D_mel_Greece_Athens_1 | KP969752 |
| LJV | D_mel_Greece_Athens_2 | KP969753 |
| LJV | D_mel_USA_Athens_GA_10 | KP969754 |
| LJV | Drosophila_Portugal_Lisbon | KP969755 |
| LJV | D_mel_USA_Athens_GA_11 | KP969756 |
| LJV | D_mel_USA_Athens_GA_12 | KP969757 |
| LJV | D_sim_USA_Athens_GA_6 | KP969758 |
| LJV | D_mel_USA_Athens_GA_13 | KP969759 |
| LJV | D_mel_Portugal_Lisbon_3 | KP969760 |
| LJV | D_sim_France_Marseille | KP96976 |
| LJV | D_sim_Greece_Athens_4 | KP969762 |
| LJV | D_mel_Ghana_Accra_2 | KP969763 |
| LJV | D_mel_USA_San_Diego_CA_3 | KP969764 |
| LJV | D_mel_Ghana_Accra_3 | KP969765 |
| LJV | D_mel_Ghana_Accra_4 | KP969766 |
| LJV | D_mel_France_Montpellier_5 | KP969767 |
| LJV | D_mel_Ghana_Accra_5 | KP969768 |
| LJV | D_mel_Ghana_Accra_6 | KP969769 |
| LJV | D_mel_Australia_Coffs_Harbour | MH384268 |
| LJV | D_mel_Australia_Hunter_Valley_1 | MH384278 |
| LJV | D_mel_Australia_Hunter_Valley_2 | MH384285 |
| LJV | D_mel_Australia_Melbourne_1 | MH384312 |
| LJV | D_mel_Australia_Melbourne_2 | MH384324 |
| LJV | D_mel_Australia_Melbourne_3 | MH384332 |
| LJV | D_mel_Australia_Melbourne_4 | MH384357 |
| LJV | D_mel_Australia_Melbourne_5 | MH384368 |
| LJV | D_mel_Australia_Melbourne_6 | MH384384 |
| LJV | Apis_mellifera_Ethiopia_Ganta_Afeshum | MT681680 |
| LJV | D_suzukii_Germany | MW556743 |
| LJV | D_sim_Australia_Raeburn_Orchards | MW976818 |
| LJV | Drosophilidae_Ghana_Accra_2 | NC027128 |
| LJV | Environmental_Sample_Slovenia | OL472185 |
| NFV | D.mel_Cambridge UK 1 | OP263083 |
| NFV | D.mel_Cambridge UK 2 | OP263082 |
| NFV | D.mel_Cambridge UK 3 | OP263084 |
| NFV | Drosophilidae | KP714070 |
| NFV | D.mel_Maryborough Australia 1 | MH384302 |
| NFV | D.mel_Maryborough Australia 2 | MH384307 |

